# The N-terminal domain of ALS-linked TDP-43 assembles without misfolding

**DOI:** 10.1101/134072

**Authors:** Phoebe S. Tsoi, Kyoungjae J. Choi, Paul G. Leonard, Antons Sizovs, Mahdi Muhammad Moosa, Kevin R. MacKenzie, Josephine C. Ferreon, Allan Chris M. Ferreon

**Author notes:** **Contributions:** J.C.F. and A.C.F. designed the experiments; P.S.T. and A.C.F. performed and analyzed the ensemble CD and fluorescence spectroscopy, and single-molecule FRET experiments. P.S.T. and J.C.F. performed and analyzed the NMR experiments. K.J.C. prepared the TDP-43 DNA constructs and performed the imaging experiments. K.J.C., J.C.F. and A.C.F. purified and prepared the TDP-43 protein samples. P.G.L. carried out the analytical SEC and sedimentation velocity experiments. A.S. and J.C.F. performed the SEC-static light scattering experiments. K.R.M., M.M.M., J.C.F. and A.C.F wrote the manuscript, with inputs from all other authors.

## Abstract

TDP-43 forms inclusions in several neurodegenerative diseases, and both its N- and C-terminal domains are implicated in this process. We show that the folded TDP-43 N-terminal domain oligomerizes under physiological conditions and propose that, in full-length TDP-43, association between folded N-terminal domains enhances the propensity of the intrinsically unfolded C-terminal domains to drive pathological aggregation.

---

Amylotropic lateral sclerosis is a disorder of motor neurons that often results in a progressive fatal muscle atrophy^1^, and cytoplasmic TAR DNA-binding protein-43 (TDP-43) inclusion bodies are the salient features of both familial and sporadic disease pathologies^2^. Studies on the mechanism of TDP-43 action have revealed a complex interplay between the *loss of cellular function* and *gain of toxic dysfunction* effects of the *TARDBP* gene product^1, 3, 4^. TDP-43 comprises an N-terminal domain (TDP-43^NTD^), two RNA recognition motif single-stranded nucleic acid binding domains, and a disordered C-terminal domain (TDP-43^CTD^)^5^. The glycine rich C-terminal domain, which harbors the vast majority of disease-linked mutations^4^, self-interacts and undergoes both liquid-liquid phase separation and aggregation^6-8^. While none of the hitherto identified pathogenic mutations reside within the N-terminal domain, recent *in vivo* studies clearly indicate a role for the N-terminal domain in pathogenic aggregation^3, 9, 10^.

The TDP-43 N-terminal domain is highly conserved across the metazoan lineage^11^; although its physiological role remains elusive, it has been reported to self-associate *in vitro*^12^ and in cells^9^. Conflicting observations have been made about TDP-43^NTD^ physical properties^5, 13, 14^. Qin *et al*^13^ report that a tagged construct of TDP-43 residues 1-102 adopts an unstable ubiquitin-like fold (∼30% unfolded at 25°C, pH 4.0 in unbuffered water) and hypothesize that this unfolded state plays a role in initiating full-length TDP-43 aggregation. Mompeán *et al*^5^ find that a tagged construct of residues 1-77 is completely folded and monomeric under similar solution conditions (25°C, pH 3.8, 3 mM sodium acetate). The contradictory findings may arise from sample conditions, construct lengths, expression tags^14^, or expression/purification methods. Here, we integrate observations from ensemble biophysical techniques and single-molecule methods to assess TDP-43^NTD^ stability and self-association, focusing on near-physiological conditions but also examining a wide pH range to attempt to reconcile divergent literature reports.

Despite reports of poor TDP-43^NTD^ *in vitro* solution properties^13^, we find refolded TDP-43^NTD^ (residues 1-80, no tags) to be readily soluble and suitable for ensemble experiments at near-physiological conditions (pH 7.5, 200 mM NaCl). Thermal denaturation of TDP-43^NTD^ (αβγ buffer, pH 7.5 ± 0.05) show clear pre-transition baselines (**Fig. 1a**), indicating that the native folded state is the major ensemble species. To probe for minor species that might escape detection in ensemble-averaged experiments, and to eliminate any self-association effects, we used single-molecule Förster resonance energy transfer (smFRET) experiments that can directly resolve minor species in a mixture^15^. We labeled TDP-43^NTD^ with donor and acceptor dyes (Alexa Fluor 488 & Alexa Fluor 594; *see* **Supplementary Methods**) at residues 2 and 81, which are close in space in the folded domain (PDB 2n4p^5^). As a single doubly-labeled protein in the dilute sample (100 pM) traverses through the femtoliter-observation volume, we record fluorescence signals from the donor and acceptor dyes. Signal intensities from individual channels are analyzed to yield FRET efficiency (*E*_FRET_) histograms. Folded, doubly labeled TDP-43^NTD^ gives high *E*_FRET_ values (**Fig. 1c**, *top panel*); decreased *E*_FRET_ implies increased distance between the dyes, from which we infer TDP-43^NTD^ unfolding and expansion. Near-zero *E*_FRET_ values (gray shaded region at left in **Fig. 1c**) arising from TDP-43^NTD^ molecules with absent, photobleached, or otherwise nonfluorescent acceptor dyes are excluded from our analysis.

**Figure 1.**
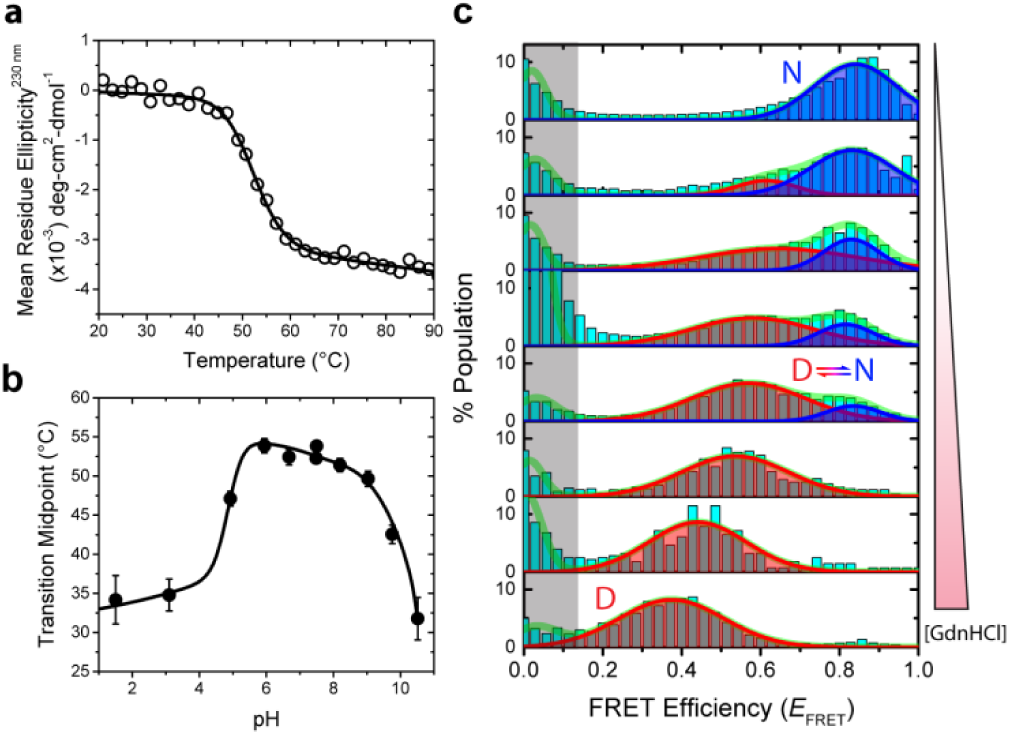
TDP-43^NTD^ is thermodynamically stable at physiological conditions. (**a**) Thermal unfolding of TDP-43^NTD^ was monitored via far-UV CD spectroscopy. Data is shown as open circles; non-linear least squares (NLS) fit of the data to a two-state protein-denaturation model is shown as a line. Transition midpoint(s) are obtained from the NLS fit. (**b**) Changes in transition midpoints as a function of pH showing TDP-43^NTD^ is thermodynamically stable at physiological conditions and access acid-and alkaline-unfolded states at respective pH conditions. *T*_m_ values are shown as filled circles; the line represents equi-population boundary of folded and unfolded states as determined through global phase diagram analysis. See also **Supplementary Fig. 7**. (**c**) Single-molecule FRET histograms of two-state unfolding of TDP-43^NTD^ upon chemical denaturation. The protein shows no detectable misfolding intermediate and at intermediate denaturant concentration undergoes slow N⇌D interconversion within the temporal resolution of our experiments (500 μs). GdmHCl concentrations for individual panels are: 0, 1, 1.25, 1.5, 1.75, 2, 4, and 6 M (top to bottom).

Binned smFRET data in the absence of denaturant are consistent with a single-distribution *E*_FRET_ value of ∼0.85, and we infer the existence of a single, folded population at pH 7.5 (**Fig. 1c**, *top*). Chemical denaturation with guanidine hydrochloride (GdnHCl) lowers *E*_FRET_ (with ∼1 M GdnHCl, *E*_FRET_ ≈ 0.65). At intermediate denaturant concentrations, histograms reveal coexisting *E*_FRET_ distributions for both folded and unfolded protein states (**Fig.** 1c, *panels 2-4 from top*), indicating slow interconversion between the states on the experimental time scale (∼500 μs). The native state mean *E*_FRET_ remains relatively unchanged from 0 to 2 M denaturant; we infer that the native state does not expand significantly with denaturant but is depopulated in favor of the unfolded state. In contrast, the denatured ensemble mean *E*_FRET_ decreases at elevated GdnHCl (**Fig. 1c** and **Supplementary Fig.13**), similar to denaturant-induced expansion of unfolded ensembles of globular proteins^16, 17^.

The TDP-43^NTD^ thermal transition midpoint for unfolding (*T*_m_) is higher than 50°C from pH 6 to 8 but drops dramatically with extremes of pH (**Fig. 1b** and **Supplementary Fig. 7**). The steep dependence of *T*_m_ on pH under acidic conditions may explain how experiments using different constructs and buffer conditions access the unfolded state to different extents. At near-physiological salt and pH conditions, however, our ensemble and single-molecule data establish that TDP-43^NTD^ is folded from low pM to μM concentrations. Successive rounds of thermal unfolding and refolding without detectable precipitation and near identical *T*_m_’s show that TDP-43^NTD^ folding is robustly reversible near physiological pH and salt.

Because our refolded TDP-43^NTD^ is stable and soluble over a broad pH range, we used NMR spectroscopy to probe its structure. ^1^H-^15^N TROSY-HSQC spectra of deuterated TDP-43^NTD^ samples (**Supplementary Figs. 10-12**) at pH 4.0 and zero salt show well-dispersed peaks with shifts similar to those reported by Mompeán et al^5^, indicating that our TDP-43^NTD^ adopts a fold similar to their construct despite lacking a His tag and having four additional C-terminal residues. However, our TDP-43^NTD^ spectra at pH 7.0 and 50 mM NaCl differ strongly from those of Mompeán et al^5^: although the amides are well-dispersed, only half of the expected resonances are visible at 25°C **(Fig. 2a** and **Supplementary Fig. 10a and 11)**. Raising the temperature to 40°C and lowering the protein concentration increases the number of peaks but also their heterogeneity (**Supplementary Figs. 11** and **12**). However, we were able to assign the backbone chemical shifts of the domain as a H_6_GB1-TDP-43^NTD^ fusion (∼20 kDa) at pH 6.8 (**Supplementary Table 1**). The backbone shifts are similar^5^ to those reported for the isolated domain at pH 4 (**Supplementary Fig. 10b**), indicating a similar fold. We propose that TDP-43^NTD^ self-association at neutral pH broadens resonances for amides positioned near the protein-protein interface, and that both low pH and N-terminal fusions decrease self-association. Resonances that are assigned in the fusion protein but absent in our spectra of the isolated domain at pH 7.0 map to a contiguous surface in TDP-43^NTD^ corresponding to the last 3 β-strands (**Fig. 2b**), supporting our proposal.

**Figure 2.**
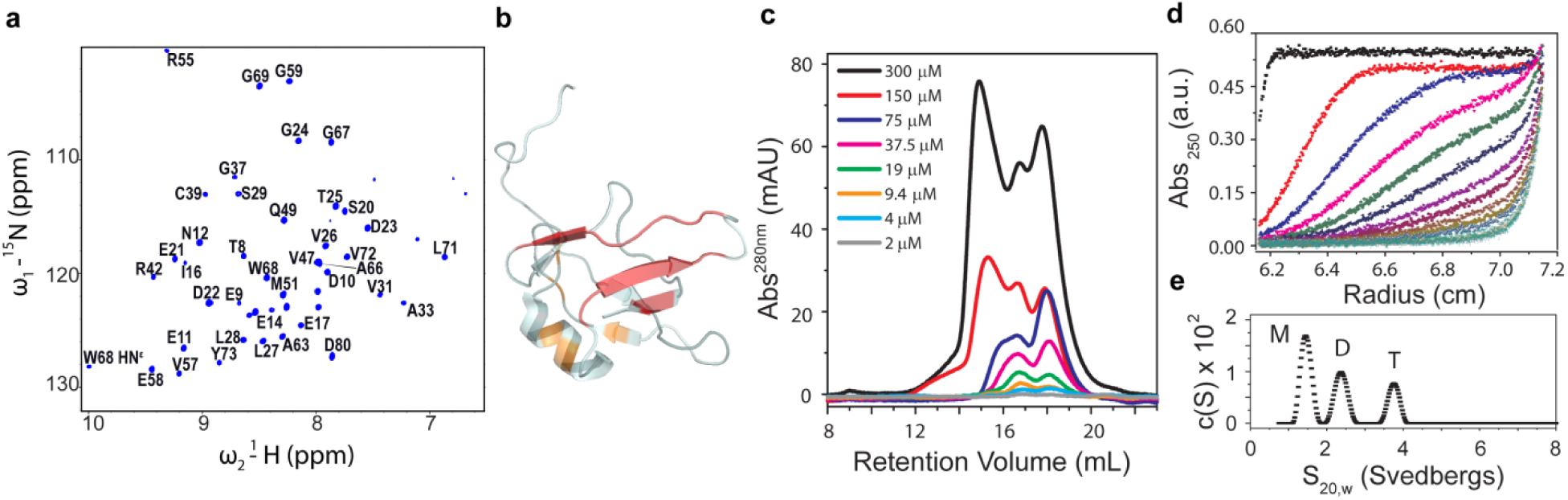
Reversible assembly of TDP-43^NTD^. (**a**) ^1^H-^15^N TROSY-HSQC spectra of TDP-43^NTD^ showing dispersed peaks, albeit with several missing resonances. The missing resonances map to a contiguous surface in TDP-43^NTD^ [(**b**), red color; PDB: 2n4p^5^]. Residues that sample alternate states are shown in orange (see **Supplementary Figure 11** for details). Size exclusion chromatography (SEC) elution profiles are shown in (**c**) for samples containing 2-300 μM TDP-43^NTD^ at physiological buffer conditions. Sequential two-fold dilution of 300 μM TDP-43^NTD^ resulted in dissociation of the high MW species (tetramer, T) to lower dimer (D) and monomeric (M) species, indicating a reversibly oligomerizing system that contrasts irreversible fibril formation ^18^. Sedimentation boundaries (**d**) and c(S) distributions (e) for 37 μM TDP-43^NTD^ from sedimentation velocity analytical ultracentrifugation experiment shows self-association in a concentration-dependent fashion (see **Supplementary Figure 9** for details).

To test directly for self-association, we performed sedimentation velocity analytical ultra-centrifugation (AUC) and size-exclusion chromatography (SEC) analyses of TDP-43^NTD^ dilution series. Both AUC and SEC show that TDP-43^NTD^ exists as monomer, dimer, and tetramer species, but not as large aggregates (**Fig. 2c**-**e and Supplementary Figs. 8 and 9**); the smallest SEC species is determined to be a monomer by static light scattering (**Supplementary Fig. 8b**). The higher order species readily dissociate to monomer and dimer, consistent with equilibrium self-association and in contrast to amyloidogenic proteins that aggregate irreversibly^18^.

Ensemble thermal unfolding fluorescence measurements show that the TDP-43^NTD^ thermal unfolding mid-point temperature (*T*_m_) increases with protein concentration **(Supplementary Figs. 2-4**). Because this finding contradicts the concentration-dependent unfolding described by Qin *et al*.^13^, we performed smFRET on doubly-labeled 100 pM TDP-43^NTD^ titrated with up to 120 μM unlabeled TDP-43^NTD^ (**Supplementary Fig. 14**). The folded state *E*_FRET_ peak is not significantly perturbed and no minor species are detected across this 10^6^-fold change in protein concentration, so we conclude that self-associating TDP-43^NTD^ maintains its global fold. Together, these data establish that the TDP-43 N-terminal domain self-associates under physiological buffer conditions through a folded, native state. The conservation of the TDP-43^NTD^ sequence^11^ suggests that oligomerization is important to TDP-43 cellular function, perhaps by modulating the affinity of the TDP-43 RNA recognition motifs for nucleic acids^12^.

Although the cellular function of TDP-43^NTD^-mediated assembly may not be fully elucidated at present, some key implications for protein dysfunction and misfolding are clear. The intrinsically disordered TDP-43 C-terminal domain (TDP-43^CTD^) is proposed to be the ‘amyloidogenic core’ that drives TDP-43 aggregation^7, 19^; at 45 μM, ∼8% of TDP-43^CTD^ can initiate assembly, resulting in phase separation^7^. Here, we have shown that TDP-43^NTD^ forms reversible oligomers to a greater extent at lower concentrations (∼30% dimer at 2-5 μM; ∼10% tetramer at 20 μM; see **Supplementary Fig. 9**). We propose that, under physiological conditions, full-length TDP-43 can assemble reversibly to dimers and tetramers via the folded TDP-43^NTD^, and that these oligomers enhance the rate of TDP-43^CTD^-mediated liquid-liquid phase separation, irreversible aggregation and/or fibril formation by raising the effective local concentration of the C-terminal domains relative to one another (**Fig. 3**). Our model explains the importance of TDP-43^NTD^ in driving a fusion protein comprising N-terminal and C-terminal TDP-43 domains flanking GFP to form giant liquid droplets with a critical concentration of phase separation of ∼5 μM^20^. The possibility that TDP-43^NTD^ itself might directly favor phase separation at physiological pH is at odds with the data presented here and is directly rebutted by its failure to do so under conditions where TDP-43^CTD^ readily phase separates and forms fibrils (**Supplementary Fig. 15**). Our model also explains why a minimal TDP-43 construct lacking RRM domains requires the TDP-43 N-terminal domain to induce aggregation of endogenous TDP-43^21^: the N-terminal domain within the artificial construct helps recruit the endogenous protein through its N-terminal domain. TDP-43 may represent a natural example of the concept demonstrated in an artificial construct by Shin et al. ^22^, where the light-inducible oligomerization domain of Cry2 dramatically accelerates liquid-liquid phase separation of a fusion protein containing a disordered domain in a light-dependent manner. Furthermore, N-terminal self-assembly may also explain why TDP-43 is found in pathological inclusions with other proteins that contain intrinsically disordered domains ^23^: if a single TDP-43 were incorporated into an aggregate via its C-terminal domain, it could recruit more TDP-43 through its N-terminal domain.

**Figure 3.**
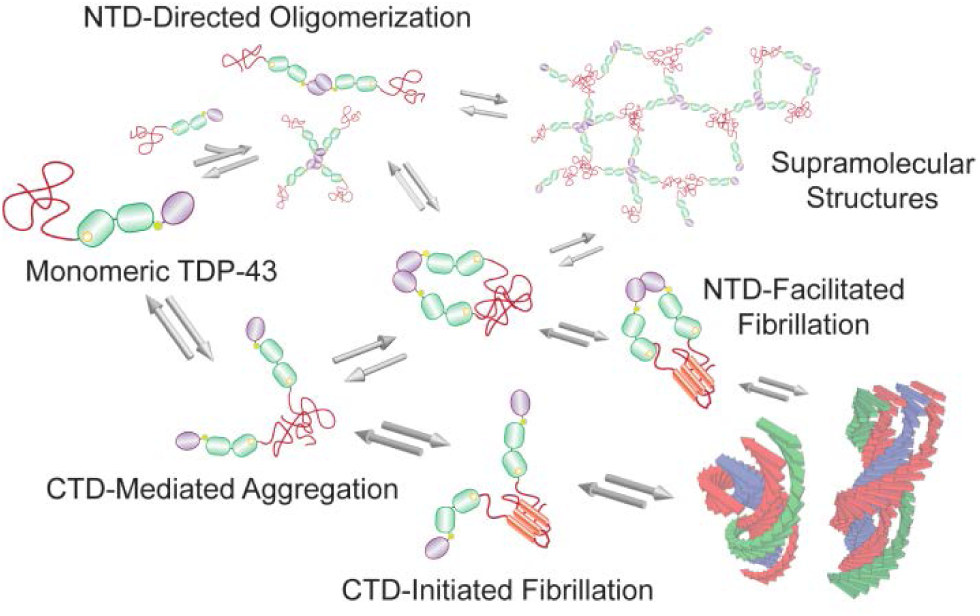
Model for TDP-43 N-terminus-facilitated C-terminal assembly and aggregation. TDP-43 comprises a fully-folded TDP-43^NTD^ (dark blue) that oligomerizes, two RNA binding domains (light green), and a C-terminal domain (TDP-43^CTD^, red) that can convert from unstructured to folded intermediate and amyloid fibril states. The N-terminus can facilitate TDP-43^CTD^ phase transitions as well as C-terminus interactions that can lead to TDP-43 fibril formation or pathological aggregation.

Other biophysical work has suggested that the unfolded state of TDP-43^NTD^ contributes to aggregation during cellular dysfunction^13, 14^. Although TDP-43^NTD^ can access the unfolded state readily at extremes of pH, we have shown that under near-physiological conditions there are no detectable misfolding intermediates by which TDP-43^NTD^ itself might aggregate. Instead, the stabe, folded N-terminal domain promotes TDP-43^CTD^-driven TDP-43 aggregation without itself misfolding, by mediating TDP-43 oligomerization.

## Acknowledgements

We thank Jin Wang’s laboratory for the use of the SEC-SLS equipment. We thank Debra Townley and Baylor College of Medicine Microscopy Core for the TEM data collection. This work was supported by laboratory startup funds from Baylor College of Medicine (A.C.F. and J.C.F.). The authors declare no competing financial interests.

